# MIRA: an open source and user-friendly software to automate counting and sizing of fungal spores

**DOI:** 10.64898/2026.08.06.743221

**Authors:** Joffrey Mejias, Henri Adreit, Anaëlle Blanc, Nadia Lubin, Cassandre Jolivet, Valentin Guyot, Oïana Brayle, Nicolas Poncelet, Elisabeth Fournier, Emmanuel Wicker, Jean Carlier, Didier Tharreau, Sébastien Ravel

## Abstract

**Background:** The quantification of fungal spores constitutes a fundamental metric in phytopathology, serving as the primary variable for inoculum standardization and being used as a proxy for disease severity. Historically, spore quantification has relied on manual hemocytometry, which remains the most precise counting process to date, where chambers such as the Malassez slide are used to count a subsample of the inoculum. However, this method applied manually is highly labor-intensive, time-consuming, and can be prone to operator-dependent variability. To overcome these limitations, we introduce MIRA (Microscopy Image Recognition & Analysis), a novel open-source software integrating You Only Look Once (YOLO) deep learning algorithms. Featuring a user-friendly graphical interface, MIRA is adaptable to multiple camera systems and supports advanced object detection models, including YOLOv11 and YOLOv26.

**Results:** We demonstrate that MIRA can be used to accurately detect and count spores from several phytopathogenic fungi, automatically measure spore surface area, and to differentiate spores across different genera. In an exhaustive comparative analysis using *Pyricularia oryzae* spores as an example, MIRA was benchmarked against manual gold-standard counting slides (Malassez and Kova) and indirect spectrophotometric methods (SPARK). The *P. oryzae* model loaded via MIRA achieved a strong correlation (R = 0.96) with manual gold standards while reducing processing time by over 90% for high-concentration samples (10⁶ spores/mL). Beyond this benchmark, we also successfully tested specific YOLO models designed to recognize macro- and microconidia of *Fusarium oxysporum f. sp. cubense*, a model for *Pseudocercospora fijiensis*, and a single multiclass model capable of identifying six different rice pathogenic fungi. We provide comprehensive tutorials for operating the software and training custom detection models for free using Roboflow and Google Colab. MIRA is available both as open-source Python code and as standalone executables for Windows and Linux.

**Conclusions:** MIRA provides a rapid, accurate, and highly reproducible alternative to manual spore counting, effectively removing a major bottleneck in phytopathology workflows. By combining advanced YOLO-based deep learning with an accessible interface and comprehensive training resources, MIRA makes accessible automated image analysis for researchers without programming expertise. Moreover, MIRA drastically improves the efficiency of high-throughput disease phenotyping and can be adapted for a wide range of microscopic quantification tasks across various biological disciplines.

## Background

In the world of plant-fungi interactions, the fungal spore is the discrete unit of infection [1]. The ability to accurately count fungal spores is essential for conducting controlled infection experiments under laboratory settings, ensuring that observed differences in disease severity are attributable to host resistance or pathogenicity rather than inoculum variability. Moreover, historically, spore morphological criteria have been the primary basis for distinguishing fungal genera or species. This expertise remains essential today, particularly for laboratories that do not have easy access to molecular biology tools. For example, in Quantitative Trait Locus (QTL) mapping for disease resistance [2,3], the intensity of sporulation on infected tissue can be a more reliable indicator of pathogen fitness and host susceptibility than visual lesion size estimation. Consequently, alongside genetic factors such as progeny size and recombination events, the statistical power of such studies remains heavily dependent on the precision of the phenotypic quantification method employed. For over a century, the standard for spore counting has been the hemocytometer (Malassez, Neubauer, Kova) [4,5]. While physically robust and inexpensive in terms of equipment, manual counting has several limitations: (i) manual counting a high-concentration sample (more than 10^6^ spores/ml) requires substantial time for a given sample, (ii) the decision to count a spore barely touching the boundaries of a grid on the hemocytometer introduces inter-operator variability, and (iii) the cognitive load of microscopic counting leads to a degradation of counting accuracy over time causing variability. Automatic counting systems have been developed using two distinct technologies, indirect quantification by spectrophotometry or the utilization of image recognition for direct quantification [6]. Indirect quantification uses turbidity as a proxy for spore density through spectrophotometry to estimate the object number based on light scattering [7]. While this method is certainly the quickest to estimate titer, it is unable to discriminate a real biological unit from background noises such as debris or mycelium. Direct quantification through image recognition has evolved from thresholding algorithms, to machine learning and more recently, deep learning [6].

Traditional computer visualization tools such as ImageJ/Fiji [8] and CellProfiler [9] remain the most widely used software in the field. These tools rely heavily on intensity-based thresholding and watershed transformations [10]. These techniques frequently fail when fungal spores exhibit internal spore structures or aggregate into dense clusters, leading to over- or under-estimations in counting. Specialized tools like OpenCFU [11] mitigate some manual labor but introduce significant bias by assuming strict circularity since it was made to count bacterial colony forming units (CFUs). Even more advanced machine learning frameworks like ilastik [12] or QuPath [13] create a workflow requiring users to navigate disjointed steps such as training, segmenting, and exporting, before a single count is generated. These limitations have historically forced researchers into a choice between the labor of manual counting and the technical barrier of image analysis.

The transition to commercial automated counters does not solve the underlying problems for plant pathology. Instruments like the Thermo Fisher Countess 3 [14] or DeNovix CellDrop [15] are fundamentally “black boxes” pre-optimized for mammalian cell lines. Their proprietary algorithms lack the transparency required for scientific reproducibility and are often incapable of being retrained for the extreme size ranges (2 µm to 100 µm) of spores from different fungal genera. While high-end systems like the µCount3D (BioSense Solutions, 2024) attempt to address these mycology-specific hurdles using 3D imaging, they introduce prohibitive capital costs and proprietary consumables.

To overcome the limitations of both manual counting and spectrophotometric assays, recent advancements in computer vision, particularly the YOLO family of object detection algorithms, have been increasingly adopted for automated, high-throughput fungal spore quantification. Several studies have demonstrated the efficacy of YOLO architectures in identifying small, dense, and morphologically complex spores across various plant pathogens while filtering out mycelial debris. For instance, Zhang et al. enhanced YOLOv5 [16] with efficient channel attention (ECA) and adaptively spatial feature fusion to detect *F. graminearum* spores, achieving over 98.5% accuracy even in complex mixed-spore suspensions. Similarly, specialized variants like MG-YOLO [17] have been engineered with multi-head self-attention mechanisms to resolve highly clustered, blurred, and multi-morphology *Botrytis cinerea* spores. To balance computational efficiency and detection accuracy for field and lab deployments, lightweight models have also emerged; the GSD-YOLO [18] model, built upon YOLOv7-tiny, incorporates decoupled heads to rapidly distinguish morphologically similar spores of *F. graminearum* and *F. asisaticum*. More recently, YOLOv8-based frameworks such as GCS-YOLOv8 [19] and YOLOv8s-SPM [20] have proven highly robust at detecting spores against natural, noisy backgrounds. However, while these advancements propose valuable algorithmic improvements and robust datasets, their outputs are predominantly restricted to raw scripts, custom network architectures, or standalone trained weights. Deploying these models in routine laboratories typically requires significant programming and bioinformatics expertise, creating a barrier for many biologists. It remains a critical need for an integrated, user-friendly platform that translates these algorithmic successes into a standardized, real-time quantification pipeline for routine laboratory and field use.To overcome these barriers, we developed MIRA (Microscopy Image Recognition and Analysis), an open-source software that automates counting on standard hemocytometer slides using deep-learning algorithms. MIRA bridges the gap between the robust detection power of deep learning and the practical needs of researchers by providing a real-time, integrated tool that ensures a 5% coefficient of variation (CV) precision without requiring coding expertise.

As a proof-of-concept of MIRA’s versatility, we developed four distinct models to detect various fungal spore genera. These include: 1) an accurate counting model for *Pyricularia oryzae*, causing blast disease on rice and wheat, 2) a classification model designed to count and discriminate between the microconidia and macroconidia of *F. oxysporum f. sp. cubense*, causing fusarium wilt of banana, 3) an instance segmentation model to count and extract the morphological dimensions of *Pseudocercospora fijiensis* spores, causing black leaf streak disease of banana and 4) a multi-class model capable of simultaneously recognizing and measuring six different genera of rice-associated fungi (*Alternaria alternata*, *F. oxysporum*, *P. oryzae*, *Bipolaris oryzae*, *Exserohilum rostratum*, and *Curvularia pisi*) within a single mixed sample. By making advanced YOLO-based architectures accessible through a simple graphical interface, MIRA provides an accurate and useful quantification solution for plant pathologists.

## Methods

### Camera and Microscope Setup

Image acquisition was performed using an Olympus BH-2 microscope equipped with a standard C-mount adapter and an Arducam 20MP IMX283 digital camera. The camera was interfaced with a Windows PC via a USB-C connection. The optical configuration consisted of a 2.5x internal camera lens coupled with either a 4x or 10x microscope objective, depending on the specific dimensions of the target fungal spores.

### Fungal Culture and Inoculation

Fungal isolates of *A. alternata*, *B. oryzae*, *E. rostratum*, *C. pisi* and *P. oryzae* were either preserved as dehydrated mycelium on filter paper and stored under vacuum at −20°C. Isolates of *P. fijiensis* and *Fusarium* sp. were stored as 15% glycerol stocks. For cultivation of fungi stored on filter paper, small fragments of the colonized filter paper were plated onto Potato Dextrose Agar (PDA: 4 g/L potato extract, 20 g/L dextrose, 15 g/L agar) and Rice Flour Agar (20 g/L rice flour, 2 g/L yeast extract, 15 g/L agar, supplemented with 1 µL/mL of a 500,000 IU/mL penicillin stock). Isolates of *Fusarium* sp. were also cultivated on PDA. These cultures were incubated for 10 days at 25°C under a standard 12-h light/12-h dark photoperiod to provide optimal conditions for vegetative growth and sporulation. For *P. fijiensis*, isolates were initially grown on PDA for 7 days. The resulting mycelium was then harvested and mechanically homogenized using a FastPrep tissue disruptor (five cycles of 20 seconds each). This mycelial homogenate was subsequently spread onto V8 juice agar plates and incubated for 12 days at 21°C under continuous illumination to induce optimal sporulation.

### Spore Harvesting and Preparation

For *P. fijiensis* spores, in each colonized V8 agar plate, 12.5 mL of sterile water supplemented with 0.02% Tween was added. The plates were then subjected to sonication to effectively detach the spores from the mycelial mat. The resulting spore suspension was subsequently collected into a sterile 50 mL conical tube (Falcon).

For other fungal species, spores were harvested after a 10-day incubation period. Five mL of osmosis water was added to each Petri dish, and the entire plate surface was mechanically scraped using a sterile L-shape spreader. The resulting crude suspension was filtered through a sterile mesh (40 µm) into a collection tube.

### Image Acquisition and Dataset Annotation

Raw microscopic images were captured at varying resolutions depending on the target fungal species and optical setup. Images of *P. fijiensis* were acquired at a resolution of 1920 × 956 pixels, while *F. oxysporum* images were captured at 2015 × 1037 pixels. Images for *P. oryzae* were captured at 3484 × 1762 pixels. The species included in the multi-class model (*A. alternata*, *B. oryzae*, *E. rostratum*, and *C. pisi*), were captured at 2910 × 1469 pixels. Image annotation and dataset management were performed using the Roboflow platform (Roboflow Inc., Des Moines, IA, USA). Depending on the specific YOLO architecture being trained, spores were manually annotated using either orthogonal bounding boxes (for object detection) or polygon masks (for instance segmentation). To ensure robust model evaluation, the global dataset for each model was partitioned into 75% training and 25% validation sets. Prior to training, standard pre-processing steps were applied within Roboflow, including auto-orientation and resizing all images to a maximum dimension of 1500×1500 pixels to match the YOLO input requirements and maximum GPU capacity. To enhance model generalization and prevent overfitting, image augmentations were applied to the training set, including random rotations, horizontal/vertical flips, brightness and exposure adjustments. The Roboflow dataset versions used to train the four models reported here are: spore-m-oryzae-xzewf v9 (https://universe.roboflow.com/cirad/m_oryzae_spore_object), spore-fusarium-object v6 (https://app.roboflow.com/cirad/spore-fusarium-object/), spore-fijiensis-seg v6 (https://app.roboflow.com/cirad/spore-fijiensis-seg/), and curvularia-genus v12 (https://app.roboflow.com/conidia/curvularia-genus/).

### Training of Deep Learning Models

All YOLO models were trained utilizing a high-performance computing environment equipped with an NVIDIA A100 80 GB GPU (MIG 4g.40gb 40 GB slice) on the ISDM-MESO HPC platform. Model weights were initialized from pre-trained Common Objects in Context (COCO) format datasets to leverage transfer learning. To ensure full transparency and reproducibility, all hyperparameters, including epochs, batch size, learning rate, and optimizer selections were saved. The exact configuration parameters and arguments utilized for training each specific fungal model are documented and available in the respective .yaml configuration files hosted on CIRAD dataverse (https://dataverse.cirad.fr/dataset.xhtml?persistentId=doi:10.18167/DVN1/E0JRDL), as well as the models.pt themselves. Training was orchestrated with the VEGA CLI (Versatile Engine for Generic Automated-training), available at https://forge.ird.fr/phim/sravel/vega, which handles Roboflow dataset download, SLURM submission, per-run reporting and canonical weight naming. Final per-model performance: *P. oryzae* (object detection): mAP@0.5 = 0.765, precision = 0.909, recall = 0.738 (best epoch 102 of 202); *F. oxysporum* micro/macroconidia (object detection): mAP@0.5 = 0.884, precision = 0.881, recall = 0.841 (best epoch 12 of 112); *P. fijiensis* (instance segmentation): Box mAP@0.5 = 0.770 / Mask mAP@0.5 = 0.693 (best epoch 72 of 196); Mix multi-genus (instance segmentation): Box mAP@0.5 = 0.823 / Mask mAP@0.5 = 0.789 (best epoch 56 of 236). Total cumulative training time on the ISDM-MESO A100 40 GB MIG slice: ∼17 h across 5 SLURM jobs.

### *P. oryzae* model performance

To evaluate counting accuracy and system performance across varying spore concentrations, six replicate spore suspensions of *P. oryzae* were prepared. Each initial suspension underwent three-fold serial dilution (yielding dilution factors of 1:4, 1:16, 1:64), resulting in a total of 24 sample tubes. To ensure sample homogeneity and experimental reproducibility, a standardized sampling protocol was implemented. Each tube was subjected to three seconds of continuous vortexing. Exactly three seconds after vortexing, the tube was held vertically, and an aliquot was extracted directly from the mid-volume of the suspension. Spore quantification for all 24 samples was performed using both Malassez and Kova counting chambers. To enable a precise comparison, a paired experimental design was utilized: each loaded slide was first assessed via traditional manual counting and subsequently analyzed using the MIRA automated model. This ensured that both counts were derived from the exact same physical sample and field of view. Concurrently, the duration of every quantification step was recorded from the moment the user or software started to count on the slide.

## Results

### Development and setup of the MIRA system

MIRA is a python-based software application featuring a Graphical User Interface (GUI) compatible with Windows and Linux operating systems. MIRA utilizes the OpenCV library for camera control (Figure 1B) and integrates the YOLO/Ultralytics deep learning framework for rapid object detection and instance segmentation. The software supports multiple YOLO architectures, including YOLOv8 [21], YOLOv11 [22], and YOLOv26 [23] (Figure 1A). Through the GUI, users can load custom model weights (.pt files) and dynamically adjust standard inference parameters. These include confidence thresholds, Intersection over Union (IoU), class labels and real-time mask visualization. MIRA also optimizes specific background processes to reduce user input. For instance, the software automatically extracts the image resizing parameters that were used during training of the model directly from the weight file and applies these exact dimensions during inference. This prevents users from having to manually keep track of and configure image dimensions for each specific model. This ensures compatibility across different models trained on varying image resolutions. MIRA is specifically designed to analyze samples loaded onto standard hemocytometers. It accommodates both Malassez and Kova slides by automatically applying their respective formulas for titer calculation (Figure 1C). Finally, the software exports the quantitative data, including spore counts, area measurements, and final calculated titers, to spreadsheet files (such as Excel) for downstream analysis. MIRA is available as a standalone executable for both Windows and Linux environments (https://dataverse.cirad.fr/dataset.xhtml?persistentId=doi:10.18167/DVN1/E0JRDL) and as open-source code via GitLab (<u>PHIM / sravel / Mira · GitLab</u>). To ensure seamless user adoption, comprehensive installation manuals and video tutorials are provided alongside the software.

**Figure 1:**
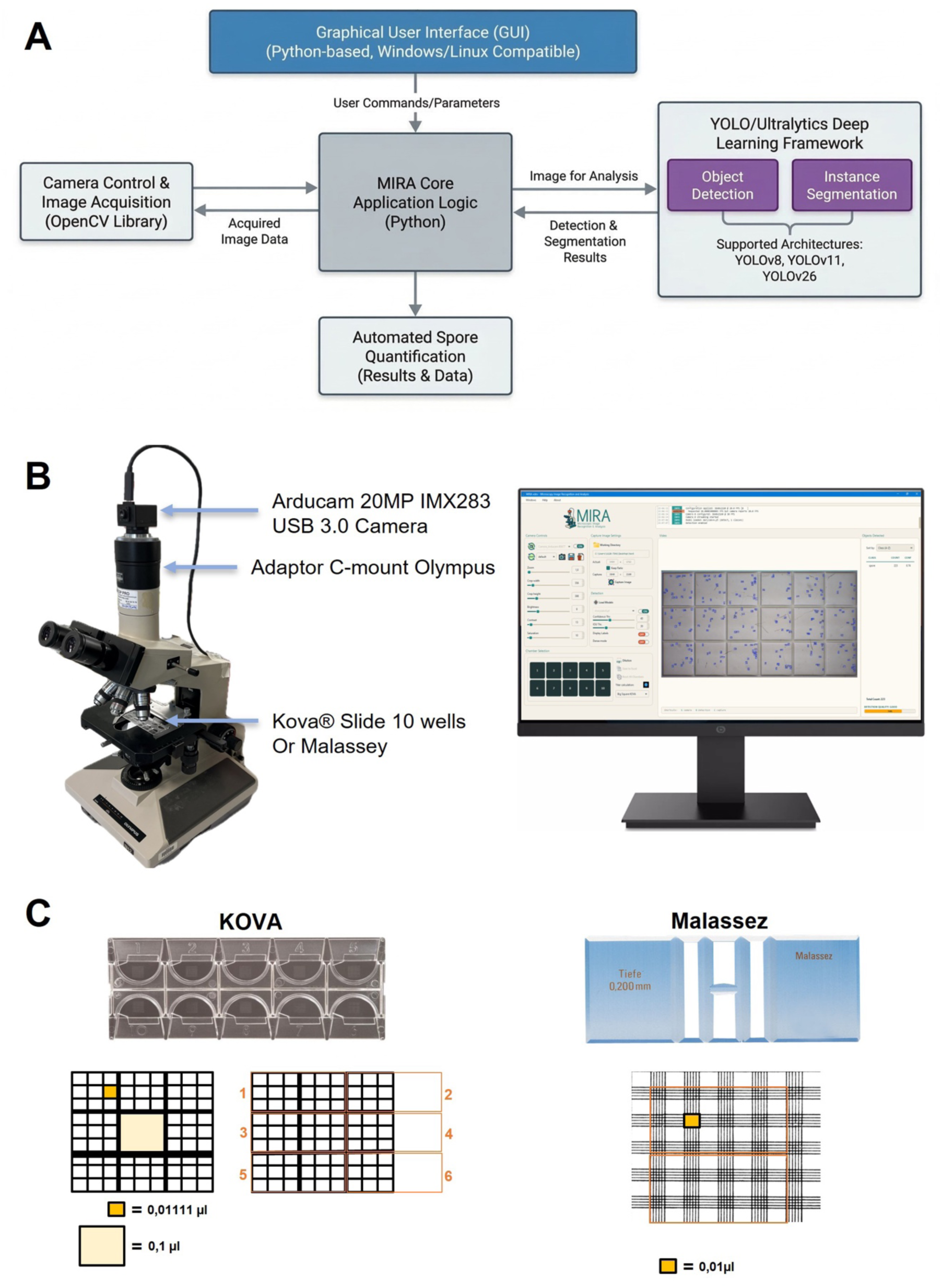
Hardware configuration and supported hemocytometer geometries for the MIRA system. (A) The standard MIRA hardware setup, featuring a standard optical microscope equipped with a C-mount adapter and a high-resolution digital camera (e.g., Arducam 20MP) to capture the slide on the stage. The MIRA Graphical User Interface (GUI) displayed on a monitor, showing real-time camera controls, volumetric calculation templates, and the live video feed with YOLO-based bounding boxes detecting spores on a hemocytometer grid. (B) On the left, a standard 10-well Kova slide. The underlying schematic details the grid layout, highlighting the volumes for a Small Square (0.0111 µL) and a Big Square (0.1 µL). The numbered sequence (1–6) illustrates the standard multi-image capture pattern required to cover the counting area. On the right, a standard Malassez counting chamber (0.200 mm depth). The corresponding schematic illustrates the grid where one small rectangle represents a volume of 0.01 µL. The orange outlines represent the software’s adjustable digital Field of View (FOV) cropping, designed to capture specific grid sections (here for 40 rectangles) for automated titer calculations.

Quantification via hemocytometry relies on a strictly defined geometric volume, for instance, a Malassez counting chamber features a grid depth of 0.200 mm and a total volume of 1 µL. The accuracy of these counts is governed by a Poisson distribution, where the coefficient of variation (CV) is inversely proportional to the square root of the total count (CV = 1/√*N*). Consequently, achieving a CV of 5% to ensure robust repeatability requires counting a minimum of 400 spores (0.05 = 1/√*N* ⇔ √*N* = 20 ⇔ *N* = 400). While reaching this threshold is highly time-consuming in a manual workflow, an automated image analysis platform with a large field of view (FOV) like MIRA can count 400 or more spores almost instantaneously (Supplemental Figure 1A).

While high-end microscope cameras often cost thousands of dollars, MIRA demonstrates that high-performance results can be achieved using affordable alternatives. In this study, we used an Arducam 20MP IMX283 USB 3.0 module (approximately 250 USD) (Figure 1). This camera mounts directly onto a standard C-mount adapter for binoculars, providing a cost-effective imaging setup capable of capturing 20 megapixels (up to 5472 × 3648 resolution) with a 60° FOV. The software automatically detects the connected camera’s profile and optimizes the resolution accordingly. For example, in our experimental setup, the software generated a live video feed (2560 × 1600 pixels) at a stable 30 frames per second (fps) for real-time visualization. Furthermore, it calibrates the count to the specific volume of the hemocytometer by allowing users to crop the detection area to an exact number of grid squares (Figure 1). This flexibility enables users to adjust the FOV based on the target object size, microscope magnification, and available sensor area.

The system is configured for image acquisition of spores in standard hemocytometers, specifically Kova slides and Malassez counting chambers (Figure 1). A Kova slide comprises 10 wells (0.9 µL total volume per well), each well is divided into 9 large squares (0.1 µL each), which are further subdivided into 9 smaller squares (0.011 µL each). A Malassez chamber holds a total volume of 1 µL, subdivided into 100 rectangular grids (0.01 µL each).

To ensure accurate and reproducible volumetric calculations, MIRA provides real-time camera controls to adjust the digital field of view (FOV) with the physical grid of the hemocytometer. To do so, users can play with the digital zoom and crop the live feed dimensions to match the specific grid layouts. The software includes standard calculation templates to accommodate various counting protocols, such as capturing two large or two small squares on a Kova slide, or analyzing 20, 40, or 100 rectangles on a Malassez chamber. Upon selection of a calculation format, MIRA automatically applies the appropriate volumetric multiplier to determine the final spore titer.

To optimize the YOLO-based detection model under varying microscopic illumination, the raw image feed can be adjusted in real-time. Modifying optical gain, brightness, contrast, and saturation ensures sufficient contrast between the target spores and the hemocytometer grid lines. For high-throughput analysis and experimental reproducibility, all user-defined parameters including spatial crops, zoom levels, and image enhancement settings, can be exported as custom JavaScript Object Notation (JSON) configuration files. These profiles can be reloaded for subsequent analyses or set as system defaults to maintain standardized conditions across different sessions and operators. Following detection and quantification (Supplemental Video 1), the resulting data can be exported directly to a .csv spreadsheet. This file contains all necessary analytical metrics, including the final calculated titer and either the average spore size, or the dimensions of each individual object, depending on the specific YOLO architecture utilized.

### Automated detection and classification of fungal spores

We evaluated if YOLO models, through the uses of MIRA, are accurate in detecting and localizing fungal spores against hemocytometer backgrounds. As demonstrated in Figure 2A, YOLO trained models can successfully apply bounding boxes to individual spores within raw microscopic images, effectively distinguishing biological targets from grid lines and background noise. In mixed samples, such as those containing both microconidia and macroconidia of *Fusarium* sp., MIRA accurately differentiated and labeled the longer, multi-septate macroconidia from the smaller, oval-shaped microconidia (Figure 2B). Detection remained highly robust even when spores were sparsely distributed or physically overlapping with the chamber’s engraved grid lines, as observed with *P. fijiensis* spores (Figure 2C). To accommodate varying image qualities and sample types, the GUI allows users to dynamically adjust the detection confidence threshold and the IoU in real-time; these parameters define the stringency of the detection and the permitted overlap allowance, respectively, enabling precise separation of adjacent or clustered objects.

**Figure 2:**
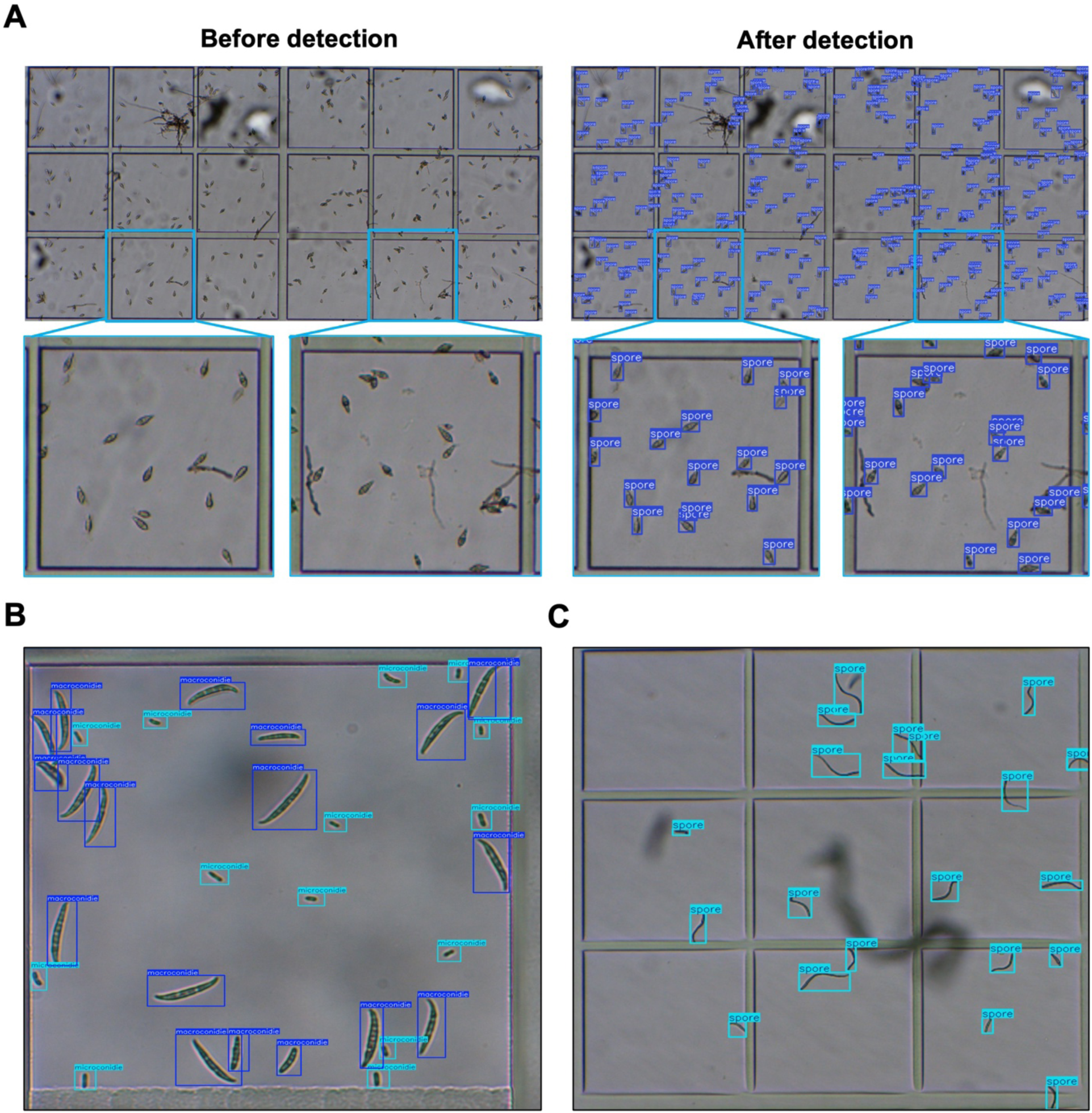
MIRA’s YOLO-based detection and classification capabilities across distinct fungal spore genera. (A) Detection of *Pyricularia oryzae* spores on a Kova chamber. The left panels display the microscopic field used in our setting to detect *P. oryzae* spores with zoomed-in regions of interest below. The right panels show the bounding boxes drawn on the screen during detection. (B) Classification of microconidia (light blue bounding boxes) and macroconidia (dark blue bounding boxes) from *Fusarium sp*. (C) Detection of *Pseudocercospora fijiensis* spores.

### Validation of quantification accuracy

To validate the counting accuracy of the YOLO models, we compared spore concentrations of *P. oryzae* (spores/mL) obtained via five different methods: Malassez manual counting, Malassez MIRA counting, Kova manual counting, Kova MIRA Counting, and a SPARK automated plate reader. For the SPARK system (TECAN, Männedorf, Switzerland), we utilized its integrated brightfield imaging module for direct object enumeration, explicitly avoiding standard indirect spectrophotometric estimations. Unlike absorbance-based methods, the SPARK imaging module captures low-resolution micrographs of the well bottom to directly identify and count discrete round shape-like objects. Counting using the different methods was performed across a four-step serial dilution (undiluted and 1:4, 1:16, and 1:64 dilutions). Across all dilution factors, MIRA automated counting yielded concentration estimates that did not significantly differ from manual counting for both Kova and Malassez slides (Figure 3A). While the Kova slides consistently yielded higher total concentration estimates than the Malassez slides across both manual and automated methods (approximately 1.5 times more), the MIRA system accurately reproduced the human-derived counts, demonstrating a near 1:1 proportional relationship. The expected logarithmic decrease in spore concentration in response to the dilution factor was observed for all methods (Figure 3A). A pooled correlation matrix further confirmed the high precision of the *P. oryzae* YOLO model on the MIRA system (Figure 4B). Linear regression analysis showed exceptional accuracy between manual and MIRA counts for the same slide types: R^2^=0.99 for the Malassez chamber (y=1.00x+4599.7) and R^2^=0.96 for the Kova chamber (y=0.96x+34969.8).

**Figure 3:**
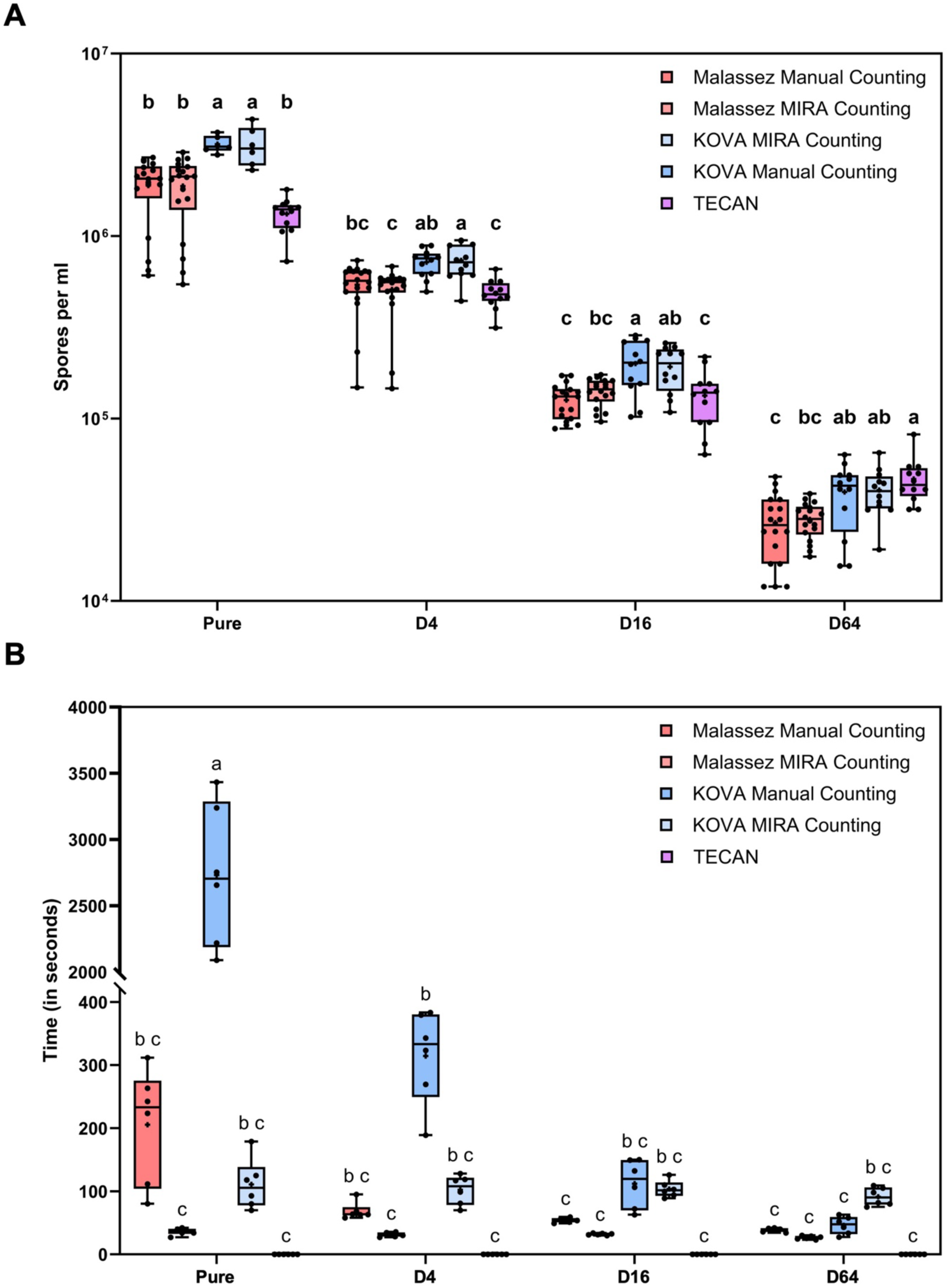
Quantitative performance and time-efficiency of MIRA compared to standard quantification methods across a dilution series. (A) Box-and-whisker plots comparing the estimated spore concentrations across undiluted and serially diluted samples (1:4, 1:16, 1:64) for the different methods employed (Manual counting, MIRA and SPARK) on two different hemocytometers (Malassez and Kova). Individual black dots represent discrete technical replicates (B) Processing time required per sample across the same dilution series. The y-axis features a scale break to accommodate the extensive time required for manual Kova counting. Box plot center lines represent the median, box limits indicate the 25th and 75th percentiles, and whiskers extend to the minimum and maximum values. Different lowercase letters (a, b, c) denote statistically significant differences between quantification methods within each dilution group (ANOVA with Tukey’s post-hoc test, *p* < 0.05).

**Figure 4:**
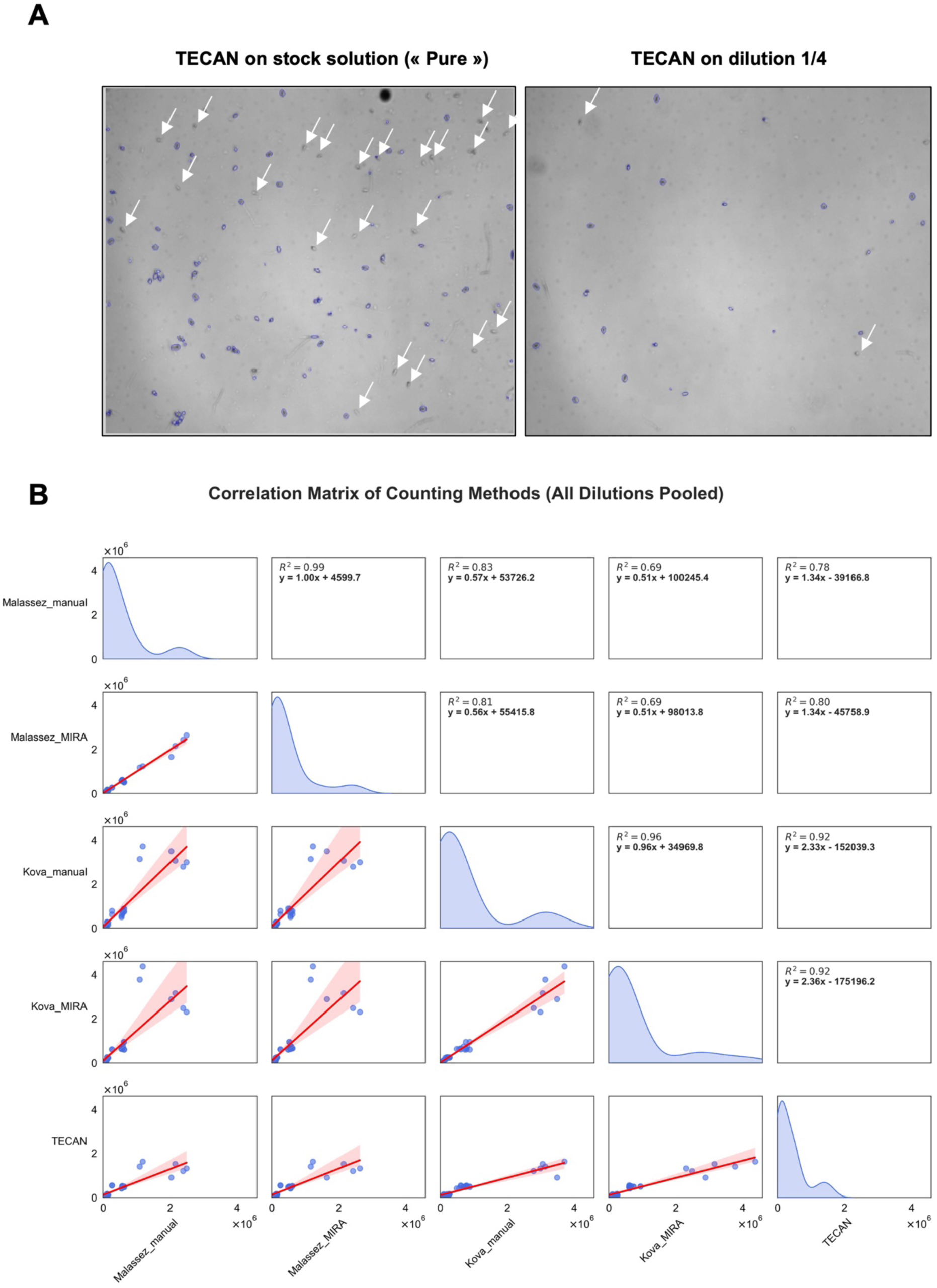
Comparative analysis of MIRA automated quantification against manual hemocytometry and optical density methods. (A) Representative microscopic fields of view illustrating spore detection by low resolution on the SPARK. (B) Correlation matrix of counting methods across all pooled sample dilutions. The diagonal plots display the data density distribution for each specific method (manual Malassez, MIRA Malassez, manual Kova, MIRA Kova, and SPARK). The lower-left panels show scatter plots comparing the methodologies, overlaid with red linear regression lines and corresponding confidence intervals (shaded regions). The upper-right panels provide the corresponding coefficient of determination and linear regression equations.

### Significant reduction in processing time

A major bottleneck in standard plant pathology workflows is the time required for manual spore quantification. To evaluate this, we recorded the time taken to count samples across a dilution series (Figure 3B). Manual counting times were highly dependent on sample concentration: undiluted samples required an average of 250 seconds for Malassez slides and up to 3,000 seconds for the larger-volume Kova slides. In contrast, MIRA’s processing time was largely independent of spore density. The automated workflow consistently required only 40 seconds for Malassez slides (capturing two images per chamber) and 100 seconds for Kova slides (capturing six images per chamber), even at the highest concentrations.

While we did not explicitly quantify processing times for extreme high-yield spore producers like *F. oxysporum*, which can yield 10⁷ to 10⁸ spores/mL (compared to 10⁶ spores/mL for *P. oryzae*), the extrapolated time savings provided by MIRA in such scenarios would be massive. Indeed, the custom detection model generated for *F. oxysporum f. sp. cubense TR4* (Figure 2B and Supplemental Figure 1) demonstrates that hundreds of spores within two small squares of a Kova slide (for a total of 81 small squares on a Kova slide) can be instantaneously counted and simultaneously classified as either macro- or microconidia.

### Comparison with standard automated optical systems

While the SPARK system exhibited near-instantaneous counting times (Figure 3B), image analysis of the SPARK captures revealed limitations in optical resolution. Images from the pure solution showed issues with spore clumping and background debris at the highest concentration, making individual spore delineation difficult compared to the spores found in the 1:4 dilution solution (Figure 4A, white arrows). Although SPARK counts correlated relatively well with Kova methods (R^2^=0.92), its correlation with Malassez methods was lower (R^2^=0.78 to 0.80). As shown in Figure 4A and 4B, spectrophotometric estimations using the SPARK reader were only comparable to manual Malassez counts at intermediate dilutions (1:4 and 1:16), indicating significantly reduced reliability at both high and low spore concentration extremes. These data suggest that while standard automated microplate readers offer rapid processing, they lack precision across a wide dynamic range. In contrast, MIRA provides a superior analytical balance by combining the high-throughput speed of automated systems with the visually verifiable accuracy inherent to traditional microscopic counting.

### Modeling multi-class spore mixture using instance segmentation

Finally, to evaluate the class multiplexing and morphometric capabilities of YOLOv26 models, we developed a comprehensive instance segmentation model (designated as the “Mix” model). This model was designed to simultaneously count, extract spore size (surface area), and perform taxonomic classification across six distinct fungal genera: *Bipolaris*, *Exserohilum*, *Curvularia*, *Pyricularia*, *Alternaria*, and *Fusarium* (Figure 5). The model was additionally trained to identify and segment generic mycelial fragments alongside the spores. We observed that the Mix model successfully discriminates between the different genera and accurately extracts the morphological dimensions of individual spores, performing robustly in both homogeneous (Figure 5A) and heterogeneous samples (Figure 5B).

**Figure 5:**
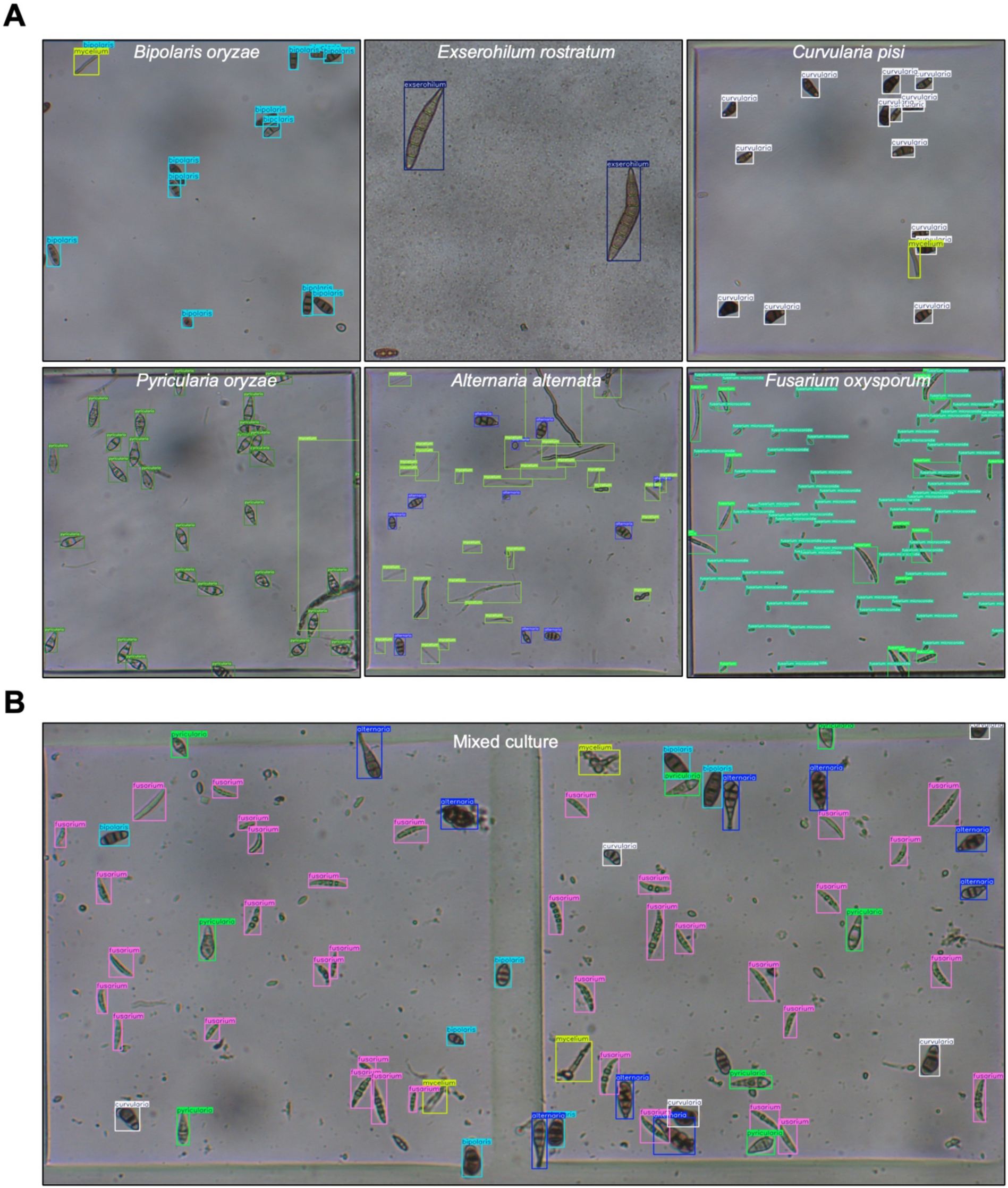
Multi-class detection and classification of six fungal spores. (A) Validation of the multi-class model on homogeneous spore suspensions. The panels demonstrate the YOLO model and software’s ability to accurately identify and apply class-specific bounding boxes to spores from distinct phytopathogenic species: *Bipolaris oryzae*, *Exserohilum rostratum*, *Curvularia pisi*, *Pyricularia oryzae*, *Alternaria alternata*, and *Fusarium oxysporum*. (B) Simultaneous multi-class detection in a heterogeneous mixed culture of the genera.

## Discussion

Accurate quantification of fungal spores is a cornerstone of experimental plant pathology, particularly when standardizing inoculum for evaluating pathogenicity/resistance or assessing disease severity directly at the sporulation level [24,25]. Precise phenotyping of fungal reproduction level is also critical for robust QTL mapping and evaluating the fitness of pathogenic mutants. However, traditional quantification methods present a significant bottleneck for large-scale phenotyping [26]. Manual hemocytometer counting is notoriously time-consuming, labor-intensive, and subject to high inter-operator variability, which can introduce detrimental statistical noise into genetic association analyses. Conversely, while automated spectrophotometric methods offer higher throughput, they critically lack the optical resolution required to distinguish true spores from mycelial fragments or background debris, often leading to bias in spore quantification [27].

A primary advantage of the MIRA system is the drastically reduced processing time without sacrificing accuracy. Our results show that YOLO models maintain a near-perfect correlation with traditional manual counting (R^2^=0.96 to 0.99, depending on the slide type) while completely decoupling the counting time from the sample’s spore concentration. For instance, the manual quantification of undiluted stock suspensions using high-volume Kova slides can require upwards of 50 minutes (around 3,000 seconds) per sample. This prohibitive timeframe often compels researchers to routinely dilute their samples to accelerate the counting process. However, this practice inadvertently amplifies the coefficient of variation by introducing pipetting errors and reducing the absolute number of counted spores, thereby compromising the statistical reliability of the final titer. MIRA reduced this to under two minutes and eliminates the subjective user bias and fatigue typically associated with prolonged manual microscopy.

Furthermore, this software circumvents the distinct limitations of standard automated microplate readers, such as the SPARK system evaluated in this study. While microplate readers provide near-instantaneous readings via optical density or low-resolution event counting, they are highly vulnerable to false positives in complex biological suspensions. Moreover, the prohibitive cost of the SPARK system and its specialized microplates limit its accessibility. Our approach circumvents this financial barrier, providing a budget-friendly alternative without compromising data accuracy. As observed in our undiluted solution analyses, microplate readers struggle with spore aggregation and background debris, leading to lower correlations with standardized manual counts (R^2^=0.78 to 0.80 for Malassez). YOLO models can identify and delineate individual spores, successfully distinguishing them from chamber grid lines, air bubbles, and host tissue debris, artifacts that routinely bias absorbance-based or simple particle-tracking algorithms.

Beyond simple enumeration, a major strength of the MIRA system lies in its capacity for multi-class analysis (object detection or instance segmentation). Many agriculturally significant fungal pathogens, such as *Fusarium* species, produce multiple spore types that play different roles in the disease cycle. The ability of MIRA to accurately differentiate and independently quantify macroconidia and microconidia within the same field of view offers a distinct advantage over flow cytometry and standard cell counters, which can struggle to separate these populations. Furthermore, because MIRA operates effectively with standard optical microscopes, commercial high-resolution cameras, and generic hemocytometers, it avoids the need for expensive, proprietary microfluidic cartridges or highly specialized imaging equipment. This democratizes the technology, allowing standard plant pathology laboratories to easily upgrade their existing equipment into AI-driven analytical tools.

We observed a systematic discrepancy between glass Malassez chambers and plastic Kova slides (Kova providing around 1.5x more spores than Malassez). Since volumetric calculation errors were ruled out, this difference is likely driven by the physical properties of the materials. The electrostatic nature of plastic Kova slides can promote spore agglutination along the grid lines, disrupting the homogeneous distribution typically seen on hydrophilic glass. Also, slight differences in capillary filling and local evaporation rates may artificially alter spore concentrations. Furthermore, the fundamental difference in loading mechanics between the two chambers is likely the primary driver of this discrepancy. A Malassez chamber is typically prepared in a manner that allows the suspension to spread uniformly beneath the coverslip, ensuring a homogeneous distribution of spores. In contrast, Kova slides are loaded via capillary action from a dedicated corner port. This directional fluid flow generates a heterogeneous concentration gradient, effectively sweeping and accumulating spores directly over the counting grid. Consequently, this leads to an artificially inflated local concentration on the Kova slide compared to the uniform distribution achieved on the Malassez chamber.

Crucially, this discrepancy does not compromise MIRA’s utility since inoculum standardization relies on relative quantification between conditions and not absolute quantification. To ensure reproducibility, researchers simply need to use one slide type consistently throughout an experiment, though we recommend the glass Malassez chamber as the gold standard if absolute quantification is needed.

## Conclusion

MIRA represents a highly reliable, time-saving tool for fungal spore quantification. By combining the verifiable accuracy of visual microscopic inspection with the speed and analytical power of deep-learning image analysis, MIRA significantly streamlines phytopathological workflows. It offers an accessible, scalable solution for high-throughput microbiology, successfully overcoming the traditional trade-off between the precision of manual counting and the labor-intensive barriers of standard microscopy.

## List of abbreviations

AI: Artificial Intelligence
ANOVA: Analysis of Variance
C-mount: Standard optical microscope camera mount standard
CFUs: Colony Forming Units
CIRAD: Centre de coopération internationale en recherche agronomique pour le développement
CLI: Command Line Interface
COCO: Common Objects in Context
CSV: Comma-Separated Values
CV: Coefficient of Variation
ECA: Efficient Channel Attention
f. sp.: Forma specialis (special form)
FOV: Field of View
FPS / fps: Frames Per Second
GCS-YOLO: Improved YOLO model variant (GhostNet, Coordinate Attention, and SimAM)
GPU: Graphics Processing Unit
GSD-YOLO: Lightweight Decoupled Wheat Scab Spore Detection Network
GUI: Graphical User Interface
HPC: High-Performance Computing
INRAE: Institut national de recherche pour l’agriculture, l’alimentation et l’environnement
IoU: Intersection over Union
IRD: Institut de recherche pour le développement
ISDM-MESO: Information Science and Data Management - Mesocentre
JSON: JavaScript Object Notation
mAP: Mean Average Precision
MG-YOLO: Multi-head self-attention and Ghost convolution YOLO variant
MIG: Multi-Instance GPU
MIRA: Microscopy Image Recognition and Analysis
MP: Megapixels
ORCID: Open Researcher and Contributor ID
PC: Personal Computer
PDA: Potato Dextrose Agar
PHIM: Plant Health Institute of Montpellier
PT / .pt: PyTorch model weights file format
QTL: Quantitative Trait Locus
SLURM: Simple Linux Utility for Resource Management
TR4: Tropical Race 4
TSARA: Transforming Food Systems and Agriculture by Research in Partnership with Africa
UMR: Unité Mixte de Recherche (Joint Research Unit)
USB / USB-C: Universal Serial Bus / Universal Serial Bus Type-C
USD: United States Dollar
V8: Brand/type of vegetable juice agar medium
VEGA: Versatile Engine for Generic Automated-training
YOLO: You Only Look Once

## Declarations

### Ethics Approval and Consent to Participate

Not applicable

### Consent for Publication

Not applicable

### Availability of data and materials

The software and code are available at <u>PHIM / sravel / Mira · GitLab</u>. All models and training configurations are available at https://dataverse.cirad.fr/dataset.xhtml?persistentId=doi:10.18167/DVN1/E0JRDL. All training datasets are available at https://universe.roboflow.com/cirad/m_oryzae_spore_object; https://app.roboflow.com/cirad/spore-fusarium-object/; https://app.roboflow.com/cirad/spore-fijiensis-seg/); https://app.roboflow.com/conidia/curvularia-genus/.

### Author Contributions

JM and SR conceived and designed the software. JM designed the experiments. JM, SR, HA, AB, CJ, NL, VG, OB, NP performed the experiments, annotated pictures, or trained the YOLO models. JM, AB, and DT analyzed the data. JM wrote the article with input from all authors. JM, JC, SR, EF, EW and DT supervised the work. JM, JC, EF and DT provided funding for the work.

### Availability of data and materials

The software and code are available at <u>PHIM / sravel / Mira · GitLab</u>. All models and training configurations are available at https://dataverse.cirad.fr/dataset.xhtml?persistentId=doi:10.18167/DVN1/E0JRDL. All training datasets are available at https://universe.roboflow.com/cirad/m_oryzae_spore_object; https://app.roboflow.com/cirad/spore-fusarium-object/; https://app.roboflow.com/cirad/spore-fijiensis-seg/ and https://app.roboflow.com/conidia/curvularia-genus/.

## Acknowledgments

We thank Melissa Bredow for critical review of the manuscript.

