## Supplementary figures and images for "MIRA: an open source and user-friendly software to automate counting and sizing of fungal spores"

### Supplemental File 1

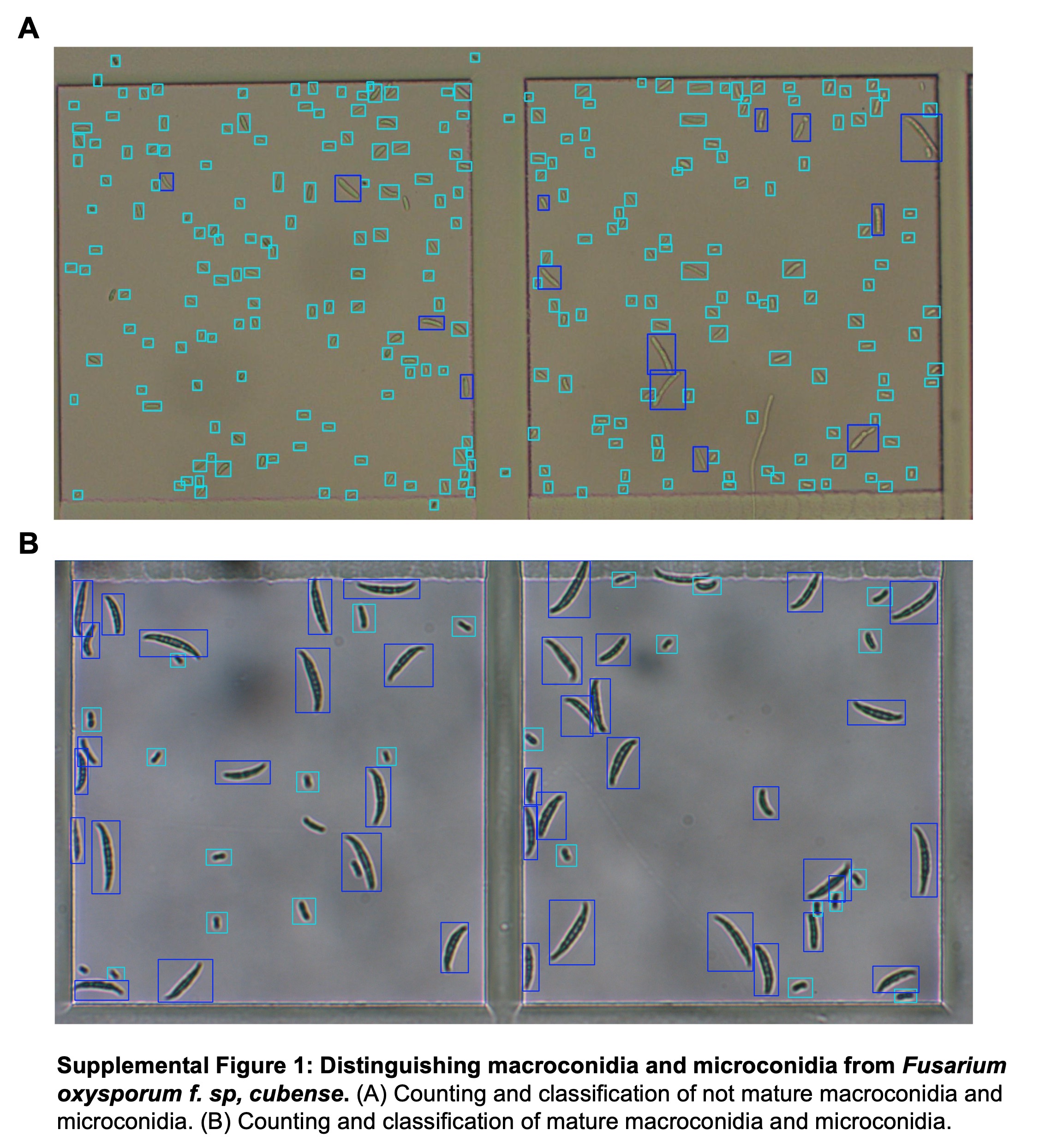
